# Sex differences in chronic social stress models in mice

**DOI:** 10.1101/605527

**Authors:** Orit Furman, Michael Tsoory, Alon Chen

## Abstract

Chronic stress creates an allostatic overload that may lead to mood disorders such as anxiety and depression. Modern causes of chronic stress in humans are mostly social in nature, relating to work and relationship stress. Research into neural and molecular mechanisms of vulnerability and resilience following chronic social stress (CSS) is ongoing and uses animal models to discover efficient prevention strategies and treatments. To date, most CSS studies have neglected the female sex and used male-focused aggression-based animal models such as chronic social defeat stress (CSDS). Accumulating evidence on sex differences suggests differences in the stress response, the prevalence of stress-related illness and the treatment response, indicating that researchers should expand CSS investigation to include female-focused protocols alongside the popular CSDS protocols. Here, we describe a novel female mouse model of CSS and a parallel modified male mouse model of CSDS in C57BL/6 mice. These new models enable the investigation of vulnerability, coping and downstream effectors mediating long-term consequences of CSS in both sexes. Our data demonstrate sex differences during CSS and for many weeks following CSS. Female mice are more prone to body weight loss during CSS and hyperactive anxious behavior following CSS. Both sexes show disturbances in social interaction, but only stressed male mice show long-term changes in neuroendocrine function and memory performance after fear conditioning. We discuss future avenues of research using these models to investigate mechanisms pertaining to sensitivity to CSS as well as treatment response profiles, in a sex-suitable manner.

## Introduction

Long-term exposure to stressful conditions is associated with the development of a manifold of pathophysiological conditions, including those affecting behavior, immune physiology, neuronal signaling, and cardiovascular function as well as chronic mood disorders such as anxiety and depression (McEwen 1998; Popoli et al. 2011; Yu 2016; Glaser and Kiecolt-Glaser 2005; Segerstrom and Miller 2004; Slavich and Irwin 2014; Boscarino and Chang 1999). In humans, chronic stress in modern life is usually social in nature, relating to socioeconomic status, work stress, relationship stress or conflicts with others e.g., social rejection or bullying (Kessler 1997, 1995; Björkqvist 2001; Slavich et al. 2010). When studying the effects of stress, it is important to examine potential sex differences, given the dramatically different effects stress has on males and females. Research in humans has established sex-differences in the biology of the HPA axis and stress response (Kudielka and Kirschbaum 2005; Bale 2006), the vulnerability to chronic stress (Hostetler and Ryabinin 2013; Bebbington 1996; Kessler and Mcleod 1984), the ensuing pathological conditions such as depression and auto-immune diseases (Whitacre 2001; Nolen-Hoeksema 1987); and finally, the response to drug treatments (Franconi et al. 2007; Kornstein et al. 2000). These data highlight the need for pre-clinical research that investigates mechanisms of pathology and resilience following chronic social stress (CSS) in both males and females.

Currently a majority of animal models of social stress, both acute and chronic, use male animals only (Tamashiro, Nguyen, and Sakai 2005). Models of psychosocial stress in rodents include manipulations of living conditions at different ages, such as maternal separation (Schmidt, Wang, and Meijer 2011), isolation stress (Hatch et al. 1963), crowding stress (Haller et al. 1999) and social instability (Hostetler and Ryabinin 2013) and resident intruder protocols, including social defeat (Koolhaas et al. 1997), maternal aggression (Lonstein and Gammie 2002) and predator exposure stress (Blanchard et al. 1998).

As many effects of psychosocial stress are sex dependent (Goel and Bale 2009), there could be important sex differences in how stress impacts behavior and physiology. Indeed, a few noteworthy research projects that used less common rodent species with unique ethological characteristics were able to describe stress-related social behaviors in both males and females, including aggression (in golden hamsters), pair-bonding (in voles) and social defeat stress (in Californian mice). In nature, Californian mice, both male and female, aggressively defend territories (Ribble and Salvioni 1990), allowing investigators to assess effects of social defeat stress in females. This sex-suitable research project revealed that the long-term effects of social defeat stress are sex-specific (Trainor et al. 2011, 2013), and has resulted in follow-up studies on cognitive flexibility following stress in mice and humans (Shields et al. 2016; Laredo et al. 2015).

The use of laboratory-reared mice allows many advantages when looking for molecular mechanisms underlying pathology, in search of novel therapeutics. The CSS protocols described in this study employ common mouse strains, and allow researchers to manipulate conditions before and after stress and its predictability while minimizing physical injuries in both males and females. Furthermore, it enables the study of short and long-term effects of CSS, and the assessment of individual differences and effects of social environment on stress response, while still offering a myriad of genetic and pharmacological tools.

In the present study, we investigated the effects of two CSS protocols on multiple measures of anxiety, depression, learning and memory, social behavior and endocrine response profile in male and female mice. Two sex-specific ethologically-minded protocols were developed in order to allow the comparative study of the long-term effects of CSS. The aims of these experiments were to obtain a clear behavioral profile after CSS and to evaluate the strengths and limitations of these models in mice.

## Materials and Methods

### Animals and Housing

Male and female (8-week old) C57BL/6 mice (Harlan) were maintained in a pathogen-free, temperature-controlled (22°C±1°C) mouse facility on a reverse 12/12 hr light/dark cycle, with lights switched on at 8 p.m. Male ICR (CD1) outbred mice (Harlan) were used as residents for the social defeat paradigm. Female ICR (CD1) outbred mice (in-house breeding) were used for the social crowding stress paradigm. All experimental animals were pair-housed with same-sex partners for the duration of the experiment. Home-cage control animals were not housed with stressed animals but with other control animals. We did not monitor females’ estrous cycle and used freely cycling females, as this factor was shown to be irrelevant for testing behavior (Prendergast, Onishi, and Zucker 2014). All experimental animals were weighed weekly to monitor general health. The Institutional Animal Care and Use Committee (IACUC) of The Weizmann Institute of Science approved all procedures. Animals were given ad libitum access to food and water.

### Female CSS paradigm

Combining elements from several social stress protocols, including resident-intruder, crowding stress and social instability, we generated a novel female mouse model of CSS. Our model employed 8-week old female C57BL/6, socially housed in groups and then in pairs in order to create highly familiar low aggression pairs.

During the 15 days of the social stress protocol, each experimental C57BL/6 female mouse was daily transported to a testing room where it was exposed to crowding by a variable number (5-9) of unfamiliar ICR female mice in a novel cage (see below description of maintaining the ICR females) for 1 h. For each stress session, the experimental mouse was allowed to interact directly with the unfamiliar ICR female mice for 30 mins; mice behavior included agonistic behavior such as sniffing and grooming as well as antagonistic behavior such as chasing, physical attacks and related vocalizations. For the remaining 30 min, the experimental mouse was placed inside a metal mesh within the same cage, enabling visual, olfactory and auditory interaction but no physical contact. In order to increase instability and unpredictability, stress sessions were conducted without habituation, each day at different times during the active dark phase and in different testing rooms. After each 1 h stress session, the experimental mice were returned to the homecage containing the same highly-familiar partner. In this manuscript, we describe behavioral and physiological response of the experimental mice to social stress, while the behavior of the partner mice is reported in a separate manuscript.

Control female C57BL/6 mice were housed in pairs and handled daily for 15 days.

The ICR female mice were housed in conditions of social instability for the duration of the social stress protocol: at the end of each stress session ICR mice were randomly divided into groups of 4 until the following day. At the beginning of each stress session, ICR mice were dispersed in novel cages containing 5-9 animals per cage. This procedure created high levels of arousal and exploration of conspecifics among ICR female mice, and rendered female ICR mice more aggressive towards the C57BL/6 female experimental mice.

### Male CSS paradigm

Based on chronic social defeat stress (CSDS) protocols in male mice (Berton, Mcclung, et al. 2006), we developed a protocol of CSDS that is comparable to the female protocol described above. Briefly, our model employed 8-week old male C57BL/6, housed in groups and then in pairs until 12 weeks old in order to create low aggression levels, as is known to occur in groups of males cohabiting for weeks (Haller et al. 1999). During the 15-day social stress protocol, each experimental C57BL/6 male mouse was transported daily to a testing room and spent 1 h in the home-cage of an aggressive and unfamiliar ICR outbred mouse (Harlan); see below the description of maintaining the ICR males. During the first 5 min of that 1 h exposure the mice were allowed to physically interact. During this time, the ICR mouse attacked the intruder mouse and the intruder displayed subordinate posturing. For the remaining 55 mins, the C57BL/6 experimental mouse was placed in a cage with a metal mesh, which enabled visual, olfactory and auditory interaction but no direct physical contact. After each 1 h stress exposure, the experimental mice were returned to their homecage, containing the same highly familiar mouse partner. In this manuscript, we describe behavioral and physiological response of the experimental mice to social stress, while the behavior of the partner mice is reported in a separate manuscript.

Control male C57BL/6 mice were housed in pairs and handled daily for 15 days.

ICR male mice were housed alone in cages for one week prior to CSDS in order to establish territoriality.

## Behavioral assessments

All behavioral assessments were performed during the dark phase following a 2 h habituation to the test room before each test. Behavioral tests were conducted in the following order, from the least stressful procedure to the most stressful: social interaction test (SIT), dark-light transfer (DLT), forced swim test (FST), fear conditioning (FC) and acoustic startle response test (ASR).

### SIT

A SIT was conducted based on prior protocols (Berton, McClung, et al. 2006; Krishnan et al. 2007b). Briefly, 24 h following the last social stress session, each experimental mouse (either male or female) was placed in a dimly illuminated open field arena (white plastic box, 50 × 50 × 22 cm, 3-5 lux) and its trajectory was tracked for two consecutive sessions of 3 min each. Video recordings were performed as previously described (Regev et al. 2011). During the first session (“no target”) the open field contained an empty wire mesh cage (circular, 10 cm diameter) located at one end of the field. During the second session (“target”), the conditions were identical except that a social target animal (an unfamiliar same-sex ICR mouse) was introduced. Between the two sessions, the experimental mouse was removed from the arena, and was placed in a temporary novel cage for approximately 1 min. An automated video-tracking software (Ethovision 9, Noldus) was used to determine the time spent by experimental mice in a defined “interaction zone” (a semi-circle surrounding the mesh) during the “No Target” and “Target” conditions. An interaction ratio was calculated as follows:

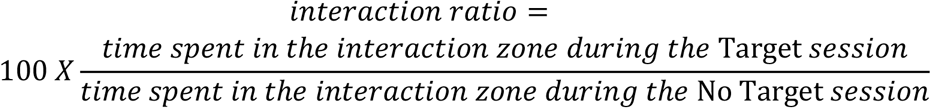

### DLT test

The DLT test takes advantage of the natural conflict of a rodent between the exploration of a novel environment and the aversive properties of a large, brightly lit open-field. The DLT test apparatus consists of a polyvinyl chloride box divided into a black dark compartment (14 × 27 × 26 cm) connected to a larger white 1200 lux illuminated light compartment (30 × 27 × 26 cm). Mice were placed in the dark chamber, and behavior was quantified during the 5 min test starting when the door connecting the dark and light chambers was opened. A video tracking system (VideoMot2; TSE Systems) was used to quantify time spent in the light compartment, the distance traveled in light area and number of dark-light transitions.

### FST

When forced to swim in a narrow space from which there is no escape, mice adopt an immobile posture after an initial period of vigorous activity, which is considered a behavioral despair reaction. The FST is considered an index of “depressive symptoms” and was performed as previously described (Issler et al. 2014). Briefly, mice were placed in Plexiglas cylinders (13 cm diameter × 24 cm high) containing water (22◦C ± 2◦C) to a depth of 10 cm, their activity during 6 minutes was recorded and immobility duration was quantified off-line with an automated video-tracking system (Ethovision 9, Noldus).

### FC

FC is a behavioral learning paradigm that allows organisms to learn to predict aversive events, by pairing an aversive event to a neutral stimulus (a foot shock paired with a neutral sounds). Previous studies reported effects of chronic stress on memory formation and retrieval using different conditioning protocols (e.g.: Rodrigues et al., 2009; Roozendaal et al., 2009), as well as sex differences in fear memory learning and extinction that were related to anxiety disorders (McDermott et al. 2015). We therefore employed a long-term protocol of FC (Karpova et al. 2011) that allowed us to examine male and female behavior during conditioning, extinction training, and long term fear memory retrieval. A computer-controlled fear-conditioning system (TSE Systems) monitored the procedure while measuring freezing behavior (defined as lack of movement except for respiration for at least 3 s); the measure was expressed as a percentage of time spent freezing. FC and extinction took place in two different contexts: FC context (A) was a transparent Plexiglas chamber cage (21 cm × 20 cm × 36 cm) with metal grids on floor and constant illumination (250 lux) whereas extinction context (B) was a black nontransparent Plexiglas chamber (same size) with planar floor and no illumination except dim red light. Both context A and context B were cleaned before each session with 70 % ethanol and 1 % acetic acid, respectively. On day 1, mice were conditioned using 5 pairings of the conditioned stimulus, CS (CS duration 30 s, 1 Hz, white noise, 80 dB) with the unconditioned stimulus, US (1 s foot-shock 0.6 mA, co-terminated with the CS); inter-trial interval (ITI): randomly varying 20-120 sec. On day 2, conditioned mice were submitted to extinction training in context B during which they received 12 presentations of the CS alone (ITI: randomly varying 30-60 s). On day 10 mice were tested first for spontaneous recovery of fear (Cue test) in context B, and two hours later were tested for context dependent fear renewal (Context test) in context A. Both tests included four presentations of the CS (ITI: randomly varying: 20-60 s).

### ASR test

The ASR test reflects a transient motor response to a sudden unexpected stimulus, such as a loud noise. Animals suffering from anxiety and hyperarousal are found to express an enhanced startle response; the startle reaction is potentiated by previous stressful experiences such as footshocks and is attenuated by prepulse inhibition (PPI: Koch 1999). Startle response and PPI protocol were assessed in a Startle Response System (TSE Systems) using a protocol adapted from Neufeld-Cohen et al. (Neufeld-Cohen et al. 2010). Briefly, mice were placed in a small Plexiglas and wire mesh cage on top of a vibration-sensitive platform in a sound-attenuated, ventilated chamber. A high-precision sensor, integrated into the measuring platform, detected movement. Two high-frequency loudspeakers inside the chamber produced all the audio stimuli. The session began with 5 min acclimation to white background noise [65 db] maintained through the whole session. Thirty-two startle stimuli [120 db, 40 ms in duration with a randomly varying ITI of 12–30 ms] were presented interspersed with an additional 40 ms startle stimuli randomly preceded by 40 ms pre-pulses of either 74 db, 78 db, or 82 dB. Latency to peak startle response and response amplitude were measured both in response to startle stimuli and in response to startle stimuli preceded by pre-pulses.

### Corticosterone measurement

Plasma was extracted from blood samples that were collected by tail bleed under basal conditions at two time points: 1 h before onset of the active phase (7 AM) and 1 h after offset of the active phase (9 PM). Blood samples were centrifuged immediately (3500 rpm for 25 min at 4°C) and extracted plasma was stored at −80°C until assayed for corticosterone (CORT) using a radioimmunoassay kit (ImmuChemTM Double Antibody Corticosterone ^125^I RIA KIT, MP biomedicals, NY, USA).

### Data analysis

Data are expressed as mean ± SEM. As stress protocols were comparable but not identical for male and female mice, we did not employ statistical models directly comparing male and female mice, but compared each stressed group to its control group. All datasets’ distributions were assessed for normality using the Kolmogorov-Smirnov test to determine which statistical tests should be employed. When comparing normally distributed data, the independent Student’s t-test was used; analysis of weight changes, reflecting within-animal comparisons necessary to address individual variability, employed Repeated Measures Analysis of Variance (RM-ANOVA), followed by post-hoc comparisons corrected by Tukey test. When data departed from normality, the Mann–Whitney U test was applied.

## Results

### Changes in body weight

As expected, both chronic stress paradigms affected body weight (BW) gain. However, female mice were affected for a longer period of time than male mice (Fig. 2 a,b). Stressed female mice (n=27) gained significantly less weight than homecage female mice (n=23) during weeks 2-6 of monitoring (including stress protocol and two weeks post-stress). RM-ANOVA revealed significant main effects for Time [weeks 2-9] [F_(7,336)_ = 150.57; P < 0.0001] and for Group [F_(1,48)_ = 12.47; P< 0.001]; the interaction Time × Group was also significant [F_(7,336)_ = 5.3; p<0.0001]. Follow-up comparisons were significant for the 3-5th weeks of weight monitoring [week 3: P< 0.0001; week 4: P<0.0001; week 5: P<0.0005; Fig. 2a]. Male stressed mice (n=13) significantly differed from homecage control mice (n=12) during week 3. RM-ANOVA indicated significant main effects for Time [weeks 2-9] [F_(7,161)_ = 147.56; P < 0.0001] but not for Group [F_(1,23)_ = 1.97; P= 0.17]; the interaction Time × Group was significant [F_(7,161)_ = 7.54; p<0.0001]. Follow-up comparisons were significant only during the 3rd week of weight monitoring [week 3: P<0.05, Fig. 2 b].

**Fig. 1.**
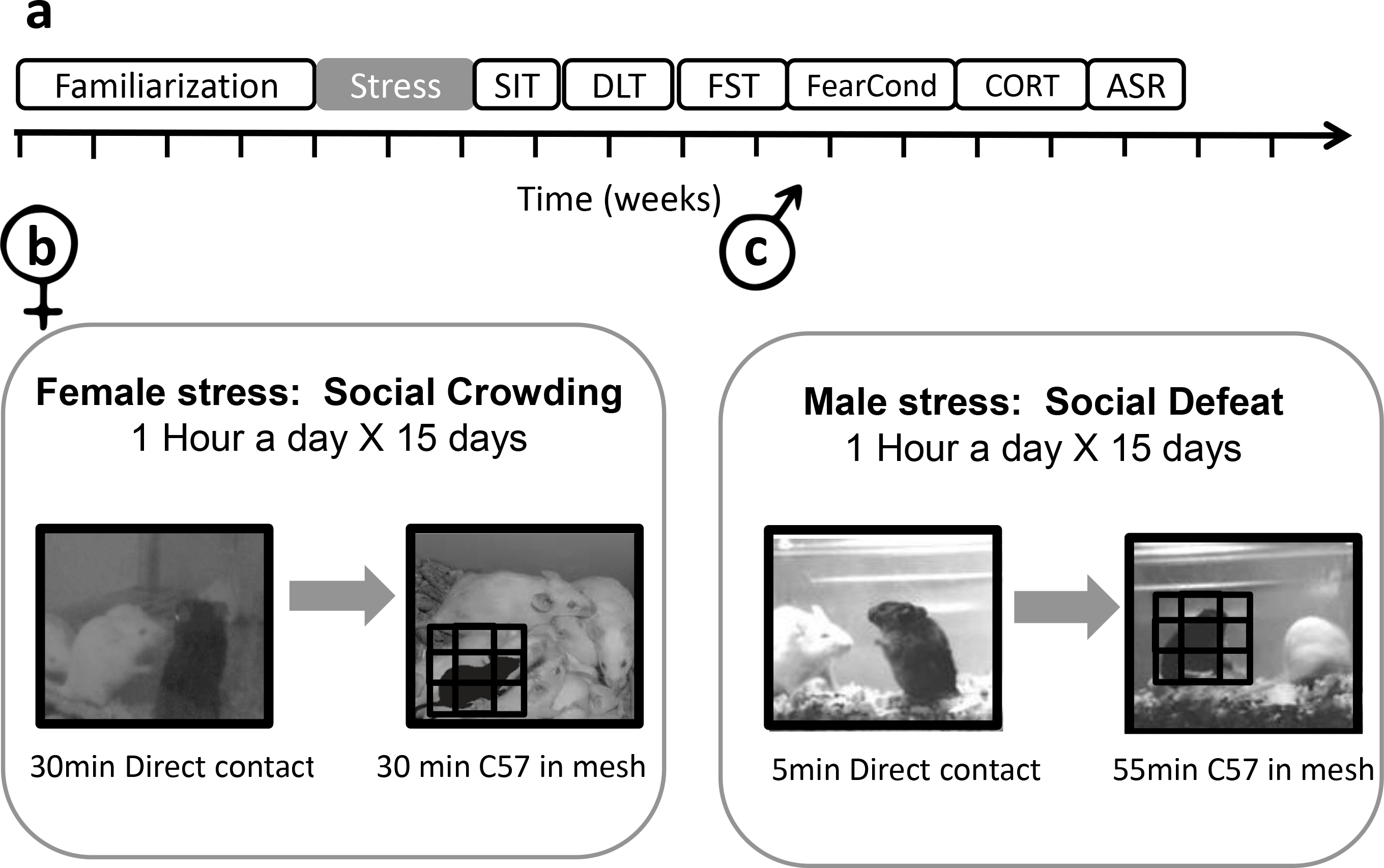
Chronic Social Stress protocols for male and female mice. **(a)** Timeline of the experimental design. **(b)** Female C57/BL mice were exposed to social crowding stress for one hour a day over 15 days, with groups of different female ICR mice. **(c)** Male C57/BL mice were exposed to social defeat stress for one hour a day over 15 days, with different male ICR mice (see *Methods* for more details). SIT – Social interaction test; DLT – Dark light transfer test; FST – Forced swim test; FearCond – Fear conditioning; CORT – Corticosterone; ASR – Acoustic startle response.

**Fig. 2.**
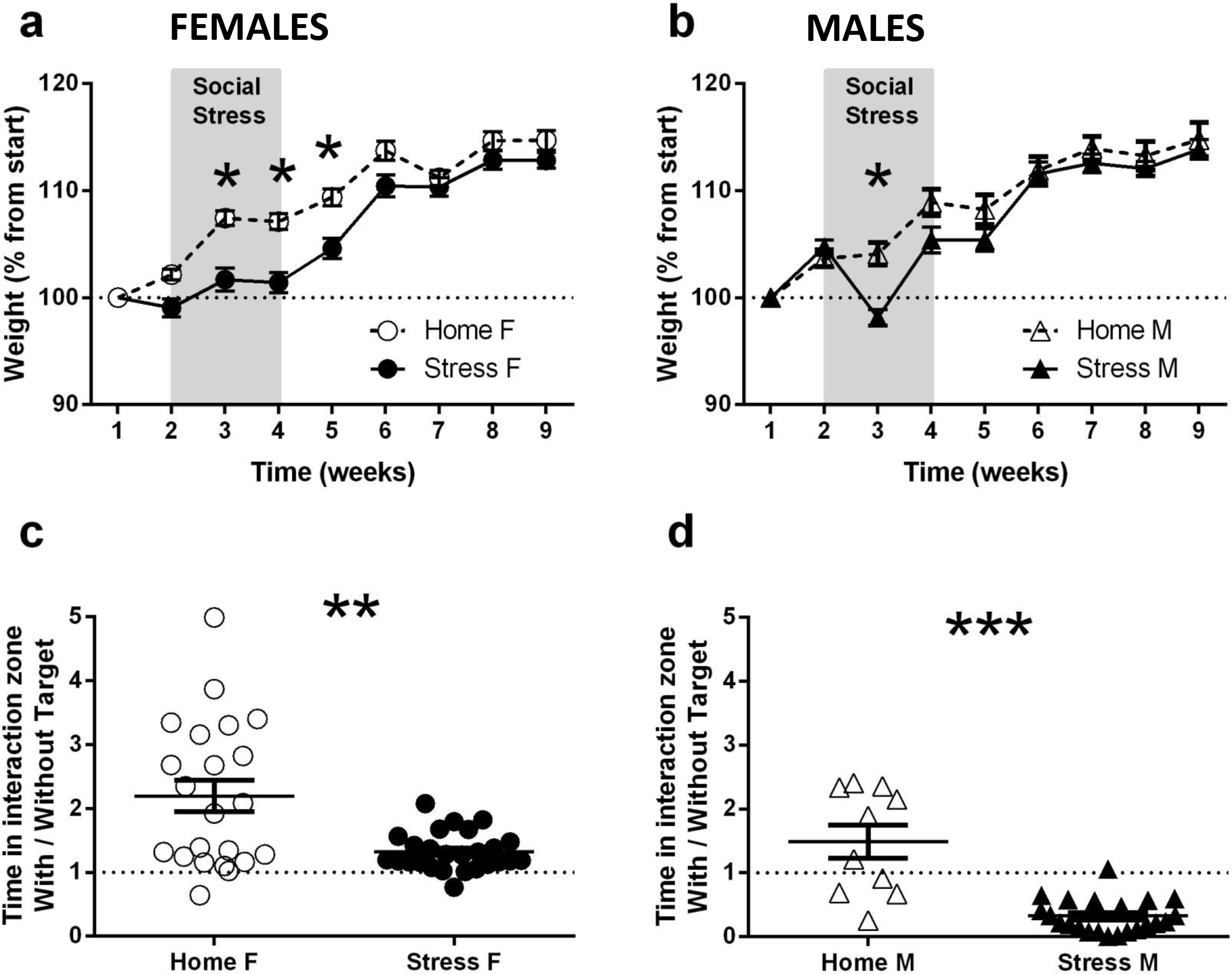
Chronic stress affected weight gain and social interaction with novel mice in both male and female mice. **(a)** Female chronically stressed mice (Stress F, n=27) gained significantly less weight than homecage control mice (Home F, n=23). Two-way repeated measure ANOVA indicated significant main effects for Time [weeks 2-9] [F_(7,336)_ = 150.57; P < 0.0001] and for Group [F_(1,48)_ = 12.47; P< 0.001]; the interaction Time × Group was significant as well [F_(7,336)_ = 5.3; p<0.0001]. Follow-up comparisons corrected by Tukey test indicated that Stress F mice gained significantly less weight than Home F mice during the 3^rd^-5^th^ weeks of weight monitoring [Denoted by an asterix, week 3: P< 0.0001; week 4: P<0.0001; week 5: P<0.0005]. **(b)** Male chronically stressed mice (Stress M, n=13) gained less weight than homecage control mice (Home M, n=12) but only during stress. Two-way repeated measures ANOVA indicated significant main effects for Time [weeks 2-9] [F_(7,161)_ = 147.56; P < 0.0001] but not for Group [F_(1,23)_ = 1.97; P= 0.17]; the interaction Time × Group was significant [F _(7,161)_ = 7.54; p<0.0001]. Follow-up comparisons indicated a significant difference only during week 3 of weight monitoring [Denoted by an asterix, week3: P<0.05]. **(c)** During the social interaction test (SIT), performed one day after end of the stress protocol, chronically stressed female mice (Stress F, n=26) show a significantly lower interaction ratio compared to female homecage control mice (Home F, n=22) (Mann-Whitney U test, U=161; P<0.01). Home F mice, but not Stress F mice, spent more time near the wire mesh when the other side contained a novel mouse vs. empty. **(d)** Similarly, during SIT performed one day after end of the stress protocol, male chronically stressed male mice (Stress M, n=22) show significantly reduced interaction ration compared to male homecage control mice (Home M, n=10) (Mann-Whitney U test, U=13; P<0.0001. Error bars represent SEM.

### SIT

It was previously shown that CSDS causes long-term reduction in social interaction, measured 24 hr after the last stress session (Tsankova et al. 2006; Berton, McClung, et al. 2006). During SIT, most control mice spent more time interacting with a social target than with an empty mesh. Therefore, we compared the interaction ratio between mice undergoing CSS (StressF, n=26; StressM, n=22) and the control group (HomeF, n=22; HomeM, n=10) and found significantly reduced interaction ratio 24 h after the last stress session in stressed mice of both sexes (Mann-Whitney U test, females U=161; P<0.01, males U=13; P<0.0001, Fig. 2c,d). An additional Chi-square test was performed on the proportion of mice categorized as “susceptible” to the stress protocol (mice with interaction scores <1), or “unsusceptible” to stress (mice with interaction scores <1). In male mice, the proportion of susceptible mice was significantly higher in the stressed vs. the control group (X^2^(1,N=32)=12.37, p<0.0005) and the vast majority of stressed mice (21 out of 22) were categorized as susceptible, as opposed to 50% in standard CSDS protocols (Krishnan et al. 2007b). In female mice, there was no difference between ratio of susceptible stressed vs. control mice (X^2^(1,N=48)=0.01, N.S.) and most stressed mice (25 out of 26) had an interaction ratio higher than 1 (Fig. 2c).

### DLT test

The DLT test took place one week after the last social stress session, in order to measure long term effects of CSS on anxiety and neophobia. We were surprised to find that stressed female mice spent more time in the light [t_(49)_ = 2.16; P<0.05, Fig. 3a], traveled greater distance in the light [t_(49)_ = 4.26; P<0.0001, Fig. 3c], made more dark-light crossings [t_(49)_ = 3.53; P<0.001, Fig. 3e] and entered the light more readily [Mann-Whitney U = 182.5; P<0.01, Fig. 3g], than female control mice. There were no significant differences between chronically stressed male mice and their respective control group [time in light: t_(33)_ = 0.75; P=0.45, Fig. 3b; distance in light: t_(33)_ = 0.8; P=0.42, Fig. 3d; dark-light crossing: t_(32)_ = 1.05; P=0.29, Fig. 3f; time to enter light: Mann-Whitney U = 90; P=0.24, Fig. 3h]. However, stressed females (Fig.3i), but not males (Fig.3j), exhibited increased average velocity during the test relative to control group mice (females: t_(49)_ = 4.9; P<0.0001; males: t_(33)_ = 1.35; P=0.18).

**Fig. 3.**
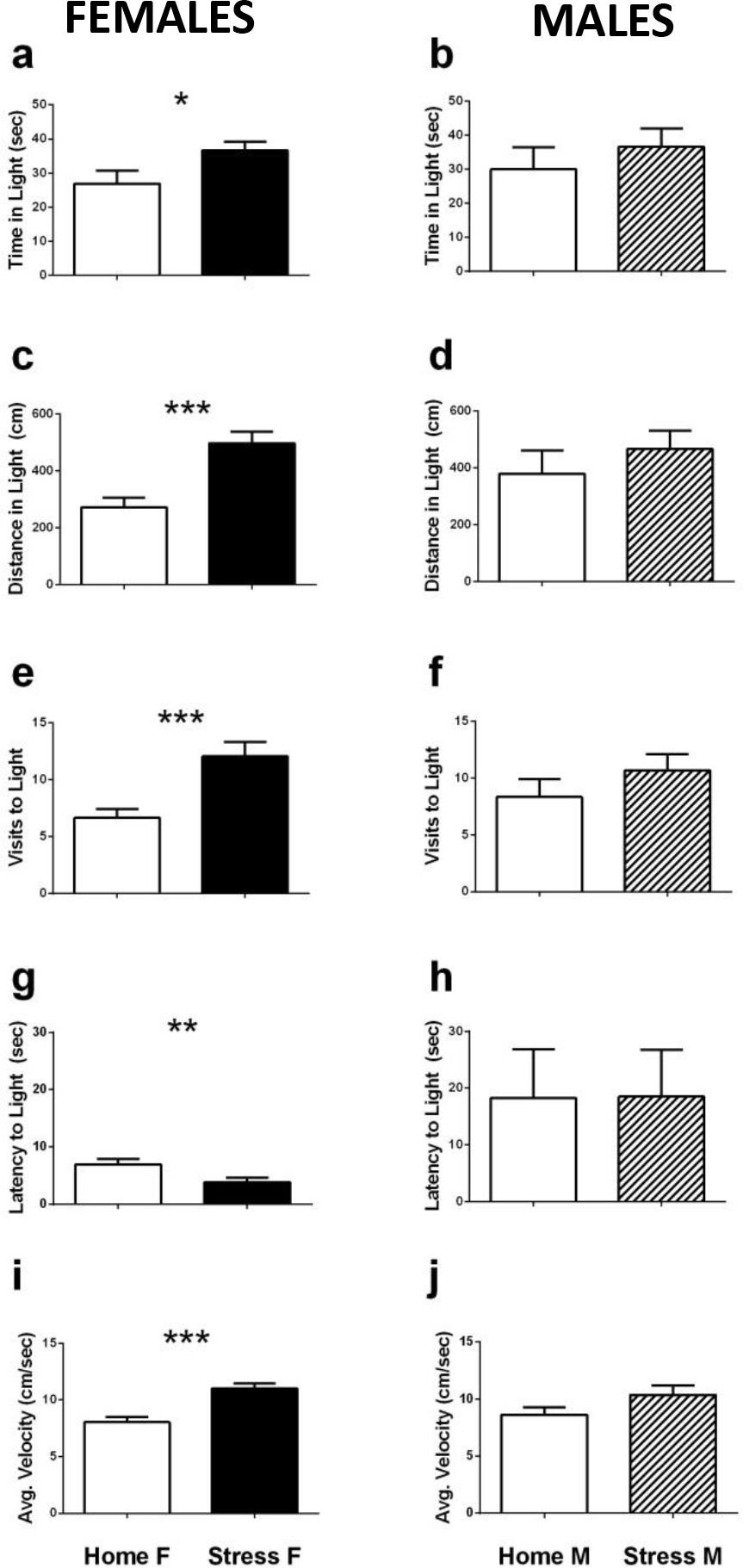
Chronic social stress affected performance in Dark Light Transfer (DLT) test of anxiety in female but not male stressed mice. Chronically stressed female mice (Stress F, n=27) spent significantly more time in light area **(a)**[t_(49)_ = 2.16; P<0.05], traveled longer distance **(c)** [t_(49)_ = 4.26; P<0.0001], made more visits to light area **(e)** [t_(43.1)_ = 3.63; P<0.005], were quicker to enter the light area **(g)**[t_(49)_ = 2.44; P<0.05] and traveled faster **(i)** [t_(49)_ = 4.9; P<0.0001] than homecage control female mice (HC F, n=24). There were no differences in any of these measures **(b,d,f,h,j)** between chronically stressed male mice (Stress M, n=23) and homecage control mice (HC M, n=12). Error bars represent SEM.

### FST

In order to measure long term effects of CSS on depression, we tested the mice in the FST 3-4 weeks after the end of the stress protocol. No differences in total immobility time were observed between the stressed and homecage control groups in either female or male mice (females: Mann-Whitney U = 232; P=0.26, Fig. S1c; males: t_(23)_ = 0.44; P=0.44, Fig. S1d). Similarly, no differences were found in latency to first immobility session in chronically stressed groups relative to their homecage controls (females: Mann-Whitney U = 270; P=0.74, Fig. S1a; males: t_(23)_ = 0.88; P=0.38, Fig S1b).

### Fear conditioning

No differences in freezing duration were observed between stress and control groups during fear conditioning and extinction (Supp. Table1). However, in the cue test, that assessed spontaneous recovery of fear, stressed males, but not stressed females, exhibited increased freezing relative to their control group (male: t_(21)_=2.09; P<0.05; female: t_(48)_=0.53; NS; Fig. 5b,c). Specifically, stressed male mice showed more freezing than controls upon initial entry into the experimental chamber, before hearing any tones, and also during later presentations of tones (Fig. 5c right panel). There were no differences in the context test that took place 2 h after the cue test, and similar freezing levels were observed between stressed and control groups, both female and male (Fig. 5 d,e, Supp. Table 1).

### Acoustic startle response test

We used the ASR test as a measure of long-term anxiety following CSS. We did not find significant differences between stressed and control groups, in neither male nor female mice, in any of the examined indices (see supplementary information and Fig. S2).

### Corticosterone

Hypothalamic-pituitary-adrenal (HPA) axis activity was assessed by measuring basal CORT levels two months after the last stress session. Only male mice showed a significant alteration in HPA activity following CSS: RM-ANOVA indicated significant main effect of group [F_(1,22)_ = 5.18; P<0.05], significant main effect of sample time [F_(1,22)_ = 46.03; P<0.0001] and non-significant interaction [F_(1,22)_ = 1.37; P=0.25]. Follow-up comparisons corrected by Fisher LSD test indicated lower levels of CORT in plasma of chronically stressed male mice, compared to controls, Before Dark [P<0.05] but not After Dark [P=0.43]. Female mice did not show significant differences beyond collection time: RM-ANOVA indicated a significant main effect of sample time [F_(1,22)_ = 48.79; p<0.0001] but non-significant group effect [F_(1,22)_ = 0.67; P=0.41] or interaction [F_(1,22)_ = 0.4; P=0.52].

## Discussion

In this study, we investigated the long-term effects of CSS in female and male mice exposed to a sex-specific ethologically valid stress protocol. A battery of behavioral tests were administered over a relatively long-time period, eight weeks following CSS, examining social interaction, anxiety and depression, learning and memory as well as the neuroendocrine profile. We found that female mice are more prone to BW loss during CSS and to hyperactive anxious behavior following CSS, both sexes show disturbances in social interaction, but only male mice show long-term changes in neuroendocrine function and in memory performance after fear conditioning.

We avoided using an identical stressor for both sexes as previous research demonstrates sex-specific vulnerability to different social stressors (Haller et al. 1999; Kessler and Mcleod 1984), and as the validity of CSDS in female lab-reared rodents is questionable, despite it being a wide-spread model for males (e.g., females are manipulated to display aggressive behavior by brain lesions). Many researchers in the field avoid exploration of sex differences related to social stress due to a lack of protocols for female rodents (Palanza 2001; Schmidt et al. 2010; Beery and Zucker 2011; Zilkha et al. 2016).

Our aim was to augment existing CSS research protocols in female mice, in order to allow meaningful exploration of sex differences in causes, effects and treatment of stress-related disorders. We therefore developed a protocol that exposed female mice to a modified crowding / social instability stress that created mild aggression between females without necessitating manipulations such as brain lesions (Haller et al. 1999), which make comparisons to males problematic. Further, we modified the CSDS protocol for males accordingly, to control for experimental parameters such as age, housing conditions, social nature of stressor, stress duration, and physical characteristics during stress exposure.

We maintained males and females under stable-paired housing conditions using a highly familiar same-sex partner throughout the experiment, as previous research has shown that social isolation is a potent stressor (Weiss et al. 2004; Hatch et al. 1963), group housing increases trait variability (Prendergast, Onishi, and Zucker 2014), and social instability is a potent social stressor in rodents (Schmidt et al. 2007). On the other hand, social support and attachment represent a major individual resilience factor for development of anxiety (Stranahan, Khalil, and Gould 2006; Isovich et al. 2001). Housing conditions were designed to preclude additional stress and to promote optimal coping with CSS. We did not monitor estrous cycle in female mice as it is an additional source of stress and as a recent extensive meta-analysis indicated neuroscientific research does not require monitoring estrous cycle in female mice (Prendergast, Onishi, and Zucker 2014).

An established physiological marker of chronic stress is a reduction BW gain. When single-sex CSDS studies were conducted, male mice and rats showed a reduction in BW gain (Krishnan et al. 2007a; Becker et al. 2008), as did female rats intruding on lactating female rats (Shimamoto et al. 2011). Two studies comparing male and female rats undergoing CSDS found BW reduction only in males (Page, Opp, and Kozachik 2016; Haller et al. 1999). We found BW reduction in both male and female mice (Fig. 2 a,b), with more pronounced and enduring effects in females and only short-term effects in males (unlike Krishnan et al. that report BW reduction in male mice several weeks after CSDS). Social instability stress was found to affect BW gain in female rats (Baranyi, Bakos, and Haller 2005) but not in female mice (Schmidt et al. 2010), though in our experiment housing conditions were stable. These findings validate the stressful nature of the CSS protocols we employed in both sexes, even when stress exposure was limited to one hour per day. Elucidation of molecular mechanisms underlying CSS effects on BW and possible sex differences awaits further investigation.

The emotional response to CSS is variable and may range between resilience and susceptibility, depending on a complex interaction between genetic and environmental elements (Nestler et al. 2002). CSDS in male mice was used successfully to describe molecular and circuit-level differences between resilient and susceptible mice, based on a behavioral phenotype of reduced social exploratory behavior, assessed by SIT (Elliott et al. 2010; Krishnan et al. 2007a). In order to assess the effect of CSS on social behavior, we employed the SIT one day after the end of CSS in both female and male mice and found a significant reduction in social interaction in both sexes (Fig. 2 c,d). When applying the published cutoff between susceptible and resilient mice at an interaction ratio of 1, we find that proportion of susceptible male mice in our modified CSDS protocol was much higher than the published 50% (Krishnan et al. 2007a). This would suggest that even 1-hour long exposure to CSDS, as opposed to 24-hour exposure in current protocols, is an extremely efficient stressor for male mice. Stressed female mice showed variability in interaction ratio but it seems that a cutoff at 1 (similar level of exploration near an empty mesh or a mesh containing an unfamiliar social target) is not appropriate to differentiate susceptible vs. resilient females, emphasizing the need to base the analysis of female behavior on empirical data obtained from female and not male subjects. Research into factors affecting resilience to stress would benefit from inclusion of females (Russo et al. 2012) as response profiles to chronic stress vary between the sexes (Beck and Luine 2002)

In order to test anxiety, we used the DLT test, a test calibrated on male rodents, where decreased activity in the light chamber is considered a marker of anxiety. We found that in several parameters, chronically stressed female mice exhibited significantly *increased* activity in the light chamber, including an *increase* in velocity. Findings of increased locomotor activity and increased exploration by female rodents during anxiety tests like DLT, elevated plus maze (EPM) and open field (OF) have previously been shown, either at baseline without previous stress exposure (Fernandes et al. 1999; An et al. 2011; Archer 1975; Johnston and File 1991), after non-social stress like restraint (R. E. Bowman et al. 2009) or inescapable shocks (Steenbergen et al. 1990). Adrenalectomy and CORT replacement did not change these findings (Kokras et al. 2012), suggesting that sex differences go beyond the baseline differences in physiology of the stress response and HPA axis function.

Several authors have suggested the motivation of male and female rodents in behavioral tests of anxiety is different, with males being driven by anxiety and females driven by activity (An et al. 2011; Fernandes et al. 1999). Archer, in a review published as early as 1975, argues that behavioral sex differences in anxiety tests showcase the different form that fearful reaction takes in different sexes, namely that males tend to become immobile and females tend to actively attempt to escape (Archer 1975). Extending these findings to behavioral differences in anxiety tests following social stressors, we find that CSS female mice are more anxious than control female mice, though this anxiety takes the form of increased locomotion, increased exploration in light and reduced latency to enter the light. CSS male mice did not differ from control male mice in this test.

We did not detect CSS related depression-like behaviors in the FST or hyperarousal and anxiety-related behaviors in the ASR or PPI. One possible reason is that FST and ASR provide behavioral readout that is not sensitive enough to detect long-term changes in behavior following CSS. It is likely that subtle long-term effects may be revealed only when readout extends to physiological measures such as heart rate, blood pressure, immune system status or brain activity monitoring during sleep (Page, Opp, and Kozachik 2016). Another explanation may be that detection of long-term effects following CSS requires anxiety-provoking social circumstances, while the tests we employed were of non-social nature. A final explanation is that the tests were conducted too long after the end of the CSS protocol, and animals had recovered from stress. However, we deem this explanation less likely, since we did find CSS-related long-term changes in memory and neuroendocrine function, as discussed below.

Learning and memory processes are affected by stress in complex ways, depending on stressor duration and intensity (acute stressors differ from chronic ones), the learning material (emotional vs. non-emotional items), the learning stage (consolidation vs. retrieval of memory), and the subjects’ age and sex (Bangasser and Shors 2010; Shors 2006; Sandi and Pinelo-Nava 2007). For example, non-social chronic stress (6 h restraint stress a day, over 21 days, in rats) was found to affect several types of memory in a sex-specific manner, with males showing impairments in object recognition and spatial memory and females showing enhanced performance or no impairment (Luine 2002; Bowman, Beck, and Luine 2003; Andreano and Cahill 2009; Bowman et al. 2009). Given this complexity, we focused on the effects of chronic stress on emotional learning using classical fear conditioning.

We investigated the effects of CSS on emotional memory in male and female mice using a protocol assessing long-term FC memory (Tsankova et al. 2006). Published research led us to speculate that this protocol may uncover non-trivial, long-term effects of chronic stress, including sex-differential effects. First, emotional memory in general, and FC memory in particular, rely on functioning and plasticity of the amygdala, hippocampus and prefrontal cortex – brain structures shown to be affected by stress, both acute and chronic (Yap et al. 2006; Roozendaal, McEwen, and Chattarji 2009; Rodrigues, LeDoux, and Sapolsky 2009). Second, retrieval of FC memory has been shown to rely on different brain structures depending on the type of retrieval: amygdala activity underlies cue memory while hippocampal activity underlies context memory (Phillips and LeDoux 1992). Third, research in humans and rodents has shown sex-dependent activity in the amygdala and hippocampus during different FC memory stages, using diverse tools such as brain imaging (Stark et al. 2006), neuronal activation (Maren, De Oca, and Fanselow 1994) and neuroanatomy (Andreano and Cahill 2009). Fourth, findings suggest that FC memory acquisition and context-triggered retrieval are enhanced in males, while females show enhanced extinction, effects that seem to depend on fluctuations in sex-hormones (Milad et al. 2010; Chang et al. 2009; McDermott et al. 2015). Fifth, social stress affects FC learning and memory as studied in socially isolated male mice and rats (Pibiri et al. 2008; Pugh et al. 1999), and one human study comparing healthy males and females suggests the effect is sex-dependent (Jackson et al. 2006).

Contrary to our literature-based expectations, we did not find stress- or sex-dependent differences during FC memory acquisition or extinction (Table S1). However, the data reveal a behavioral difference during retrieval only in male CSS mice, such that freezing was enhanced during cue-triggered memory retrieval (hearing CS in a context different than the conditioning context) but not during context-triggered retrieval (Fig. 4). This finding suggests that future research making use of the CSS protocols should focus on neurocircuitry and neurochemistry of the amygdala. Relevant research questions should pertain to long-term CSS effects on long-term emotional memory, why females seem to be resilient to enhanced retrieval compared to males, with special emphasis in assessing social factors (e.g., housing conditions, social parameters during memory retrieval) and their effects at different times after emotional memory formation.

**Fig. 4.**
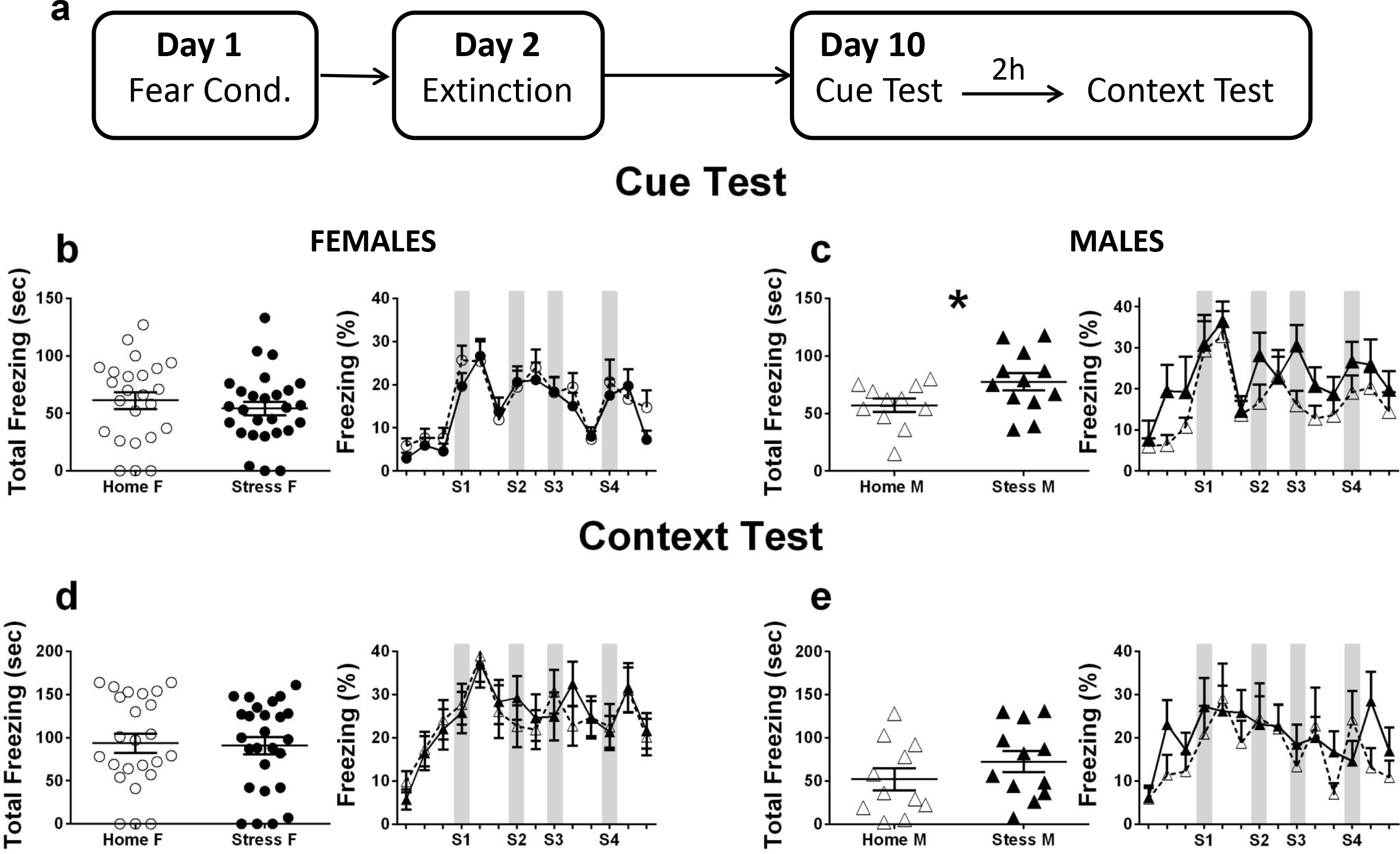
Chronically stressed male, but not female, mice show enhanced spontaneous recovery of fear 10 days after conditioning. **(a)** Timeline of fear conditioning protocol. See *Methods* for details. Panels **b-e** present behavior during 5-min cue and context tests, both administered in succession, 10 days after fear conditioning. Behavior is presented as total freezing time (left panel) and as %freezing per 30-sec bins (right panel, gray squares indicate timing of sound reminders). During the cue test, male stressed mice (**c**, full triangles on left panel, full lines on right panel, Stress M; n=12) showed significantly more freezing than homecage control mice (**c**, empty triangles on left panel, dashed lines on right panel, Home M; n=11), [t(21)=2.09;P<0.05]. There was no difference in freezing during the cue test between female stressed mice (**b**, full circles on left panel, full lines on right panel, Stress F; n=26) and homecage control mice (**b**, empty circles on left panel, dashed lines on right panel, Home F; n=24). During the context test there were no differences in freezing between chronically stressed and control female **(d)** or male **(e)** mice. Error bars represent SEM.

One of the explanations for sex differences in susceptibility to stress-induced affective disorders such as depression and anxiety is that females show heightened sensitivity to stress at multiple levels of HPA axis function. For instance, female rodents’ baseline levels of circulating CORT and corticotropin-releasing factor (CRF) mRNA expression in the paraventricular hypothalamic nucleus are higher than in males (Solomon and Herman 2009; Dunčko et al. 2001; Kitay 1961). The CRF-receptor type 1 (CRFR1) in female brains shows increased cellular signaling and decreased internalization following stress (Valentino, Van Bockstaele, and Bangasser 2013). The CRF-binding protein (CRF-BP) levels are higher in female mice (Speert, McClennen, and Seasholtz 2002) and genetic knock-out of CRF-BP affects female but not male aggression levels (Gammie, Seasholtz, and Stevenson 2008). Furthermore, negative feedback by glucocorticoids (GCs) in the brain is slower and possibly decreased due to lower glucocorticoid receptor (GR) content and less GR translocation into the nucleus in female brains (Bangasser and Valentino 2014). Finally, gonadal hormone levels were shown to affect some of these differences (CRF, CRF-BP levels, GR translocation) but not all of them (CRFR1 activity) (Bangasser et al. 2010).

Further, social environmental conditions such as group housing/crowding have been shown to affect males and females differently: isolated female rats have higher CORT level than crowded or group housed female rats whereas male rats show an opposite pattern (Brown and Grunberg 1995). In Californian mice, where male and female mice undergo CSDS, immediately after defeat, females show higher CORT than their control group but males do not (Trainor et al. 2010). However, 4 weeks after defeat, males show higher CORT levels than their control group but females do not (Trainor et al. 2011). As above, the effects of sex hormones on HPA axis activity was revealed when gonadectomy (GD) was performed before CSDS, and had an opposite effect on male and female Californian mice. GD on male mice exposed stress effects on CORT levels (no differences between stress and control group unless mice underwent GD) but GD on female mice erased stress effects on CORT (higher CORT in stressed vs. control intact females, no differences after GD; Trainor et al. 2013).

When we assessed baseline CORT levels eight weeks after the last CSS session, we found a decrease in CORT levels in males but not females (Fig. 5). This finding is in line with findings of male-specific CORT differences 4 weeks after CSDS in Californian mice (Trainor et al. 2011), though we found decreased and not enhanced CORT levels in CSS males. The difference may stem from stress resilience differences that were not assessed in the Californian mice. When examining male mice 8wks after defeat, Krishnan et al. found a decrease in AM CORT levels in susceptible mice but an increase in unsusceptible mice (Krishnan et al. 2007a). We assessed behavioral measures of resilience using the SIT and found almost all male mice were susceptible to the CSS protocol, as indicated by an interaction ratio < 1 (Fig. 2d). Resilience assessment in female mice likely requires a different cutoff point and future studies will be required to strengthen this finding and further examine sex-specific differences in stress-related CORT levels, e.g., after FST or restraint challenges.

**Fig. 5.**
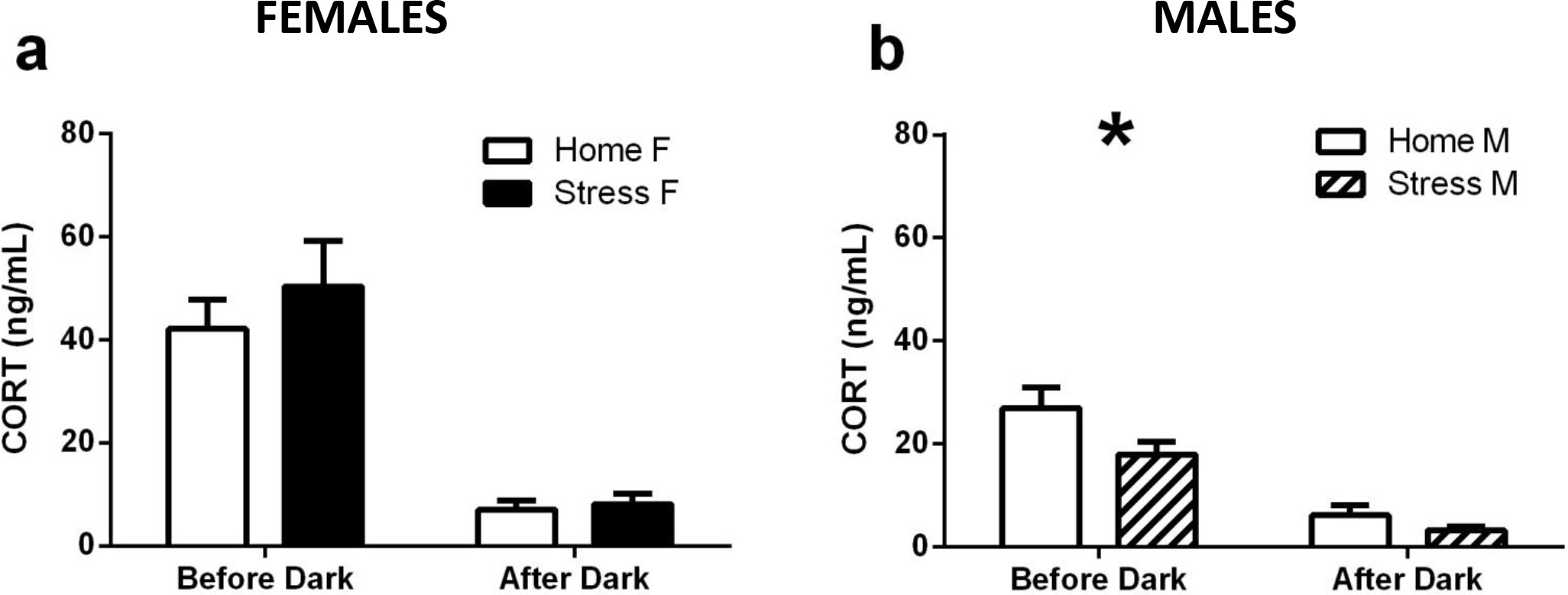
Plasma corticosterone levels were lower in male but not female stressed mice 2 months after chronic stress. Samples from tail blood were collected under basal conditions, one hour before onset of active phase (Before Dark, 7am) and one hour after the end of the active phase (After Dark, 9pm) in male and female mice. **(a)** Female mice (Home F, n=11; Stress F, n=13) showed a trend for higher CORT levels Before Dark that did not reach significance. Two-way repeated measures ANOVA indicated a significant main effect of sample time [F(1,22)=48.79; p<0.0001] but non-significant group effect [F(1,22)=0.67;NS] or interaction [F(1,22)=0.4;NS]. **(b)** Chronically stressed male mice (Stress M, n=12) showed a significantly lower CORT levels than homecage controls (Home M, n=12). Two-way repeated measures ANOVA indicated significant main effect of group [F(1,22)=5.18;P<0.05], significant main effect of sample time [F(1,22)=46.03;P<0.0001] and non-significant interaction [F(1,22)=1.37; NS]. Follow-up comparisons indicated a significant difference Before Dark [P<0.05] but not After Dark. Error bars represent SEM.

Finally, an individuals’ genetic and epigenetic background has an important role in determining their reaction to stressful environmental conditions (Schmidt, Sterlemann, and Müller 2008), and as our data indicated, there is wide phenotypic variability even among genetically identical lab-reared mice (e.g., Fig.4, Fig. S2). The CSS protocols described here could aid researchers to investigate molecular and cellular mechanisms of resilience and vulnerability to CSS in a sex-appropriate manner. Our protocols can be combined with established early-life perturbations such as maternal separation or trauma exposure during adolescence. Hopefully such combined methods will enable a better understanding of the mechanisms underlying the sex-specific susceptibility to psychopathology during adulthood that are seen in epidemiological surveys. Combining physiological readout methods as suggested above with pharmacological therapeutic interventions, such as SSRI administration after CSS, would allow investigation of sex-appropriate treatment of psychopathologies. This could create empirical support for sex-specific and perhaps individual-specific treatment regiments that will promote the wellbeing of patients that are currently suffering as non-responders to existing treatment.

## Acknowledgments

We thank Mariana Schroeder, Yael Kuperman, Ilana Adler and Maya Lebow for their contribution to this work via insightful discussions and technical expertise. A.C. is the head of the Max Planck Society–Weizmann Institute of Science Laboratory for Experimental Neuropsychiatry and Behavioral Neurogenetics. This work is supported by: an FP7 Grant from the European Research Council (260463, A.C.); a research grant from the Israel Science Foundation (1565/15, A.C.); the ERANET Program, supported by the Chief Scientist Office of the Israeli Ministry of Health (3-11389, A.C.); the project was funded by the Federal Ministry of Education and Research under the funding code 01KU1501A (A.C.); research support from Roberto and Renata Ruhman (A.C.); research support from Bruno and Simone Licht; I-CORE Program of the Planning and Budgeting Committee and The Israel Science Foundation (grant no. 1916/12 to A.C.); the Nella and Leon Benoziyo Center for Neurological Diseases (A.C.); the Henry Chanoch Krenter Institute for Biomedical Imaging and Genomics (A.C.); the Perlman Family Foundation, founded by Louis L. and Anita M. Perlman (A.C.); the Adelis Foundation (A.C.); the Marc Besen and the Pratt Foundation (A.C.); and the Irving I. Moskowitz Foundation (A.C.).

## Author Contributions

O.F. conceived and designed the experiments, collected the data, performed data interpretation and analysis, drafted the paper and approved the final version of the paper to be published. M.T. assisted in collecting the data, data interpretation and analysis, provided critical revision of the paper and approved the final version of the paper to be published. A.C. provided guidance in conceiving and designing the experiments, provided critical revision of the paper and approved the final version of the paper to be published.

---

**Fig. S1.**
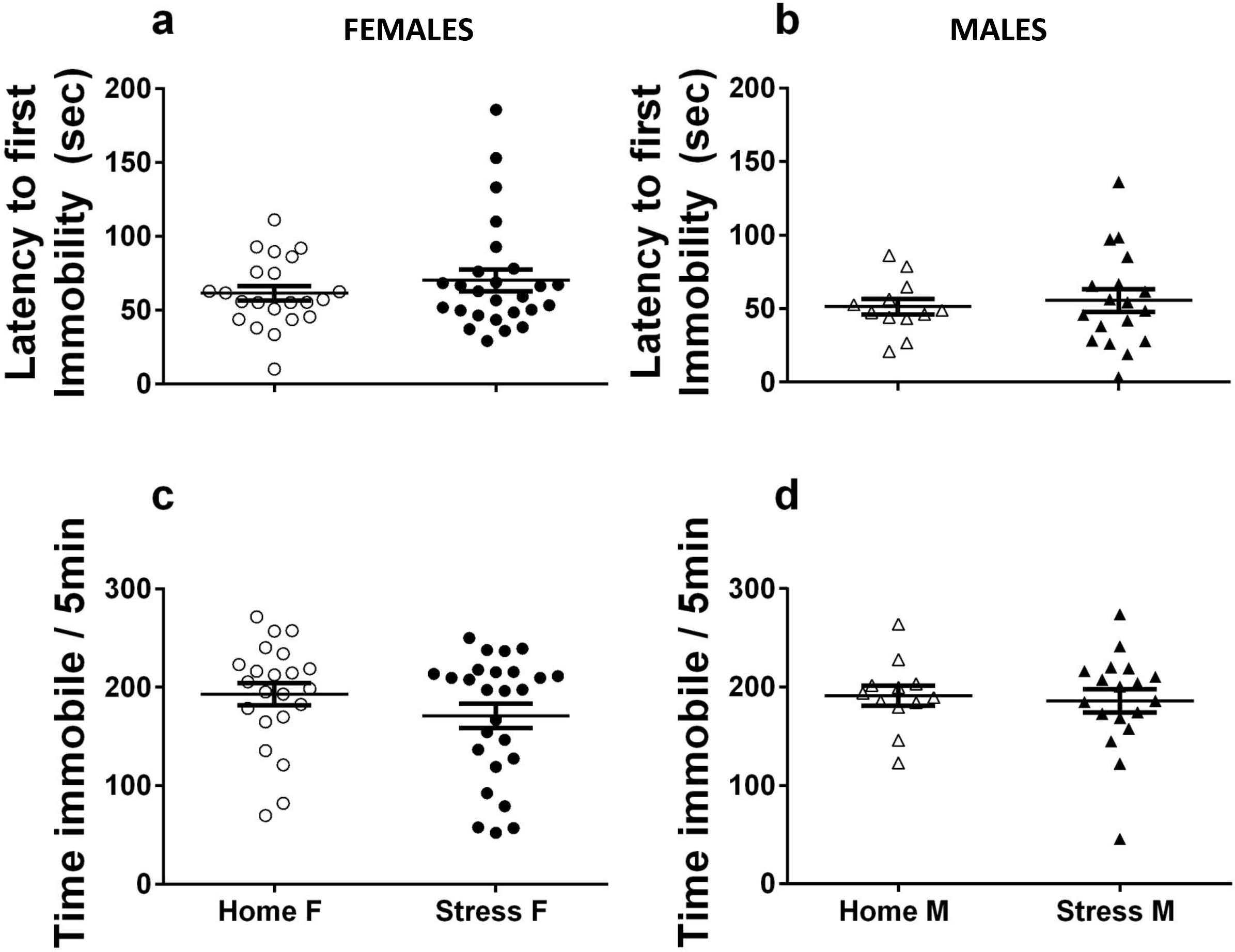
Chronic stress did not affect freezing during Forced Swim Test in male or female mice. Both female and male mice showed no difference in latency to first period of freezing **(a,b)**, or in total duration of freezing during the last five min of a six min swim session **(c,d).** (Home F: n=22; Stress F: n=26; Home M: n=12; Stress M: n=18). Error bars represent SEM.

**Fig. S2.**
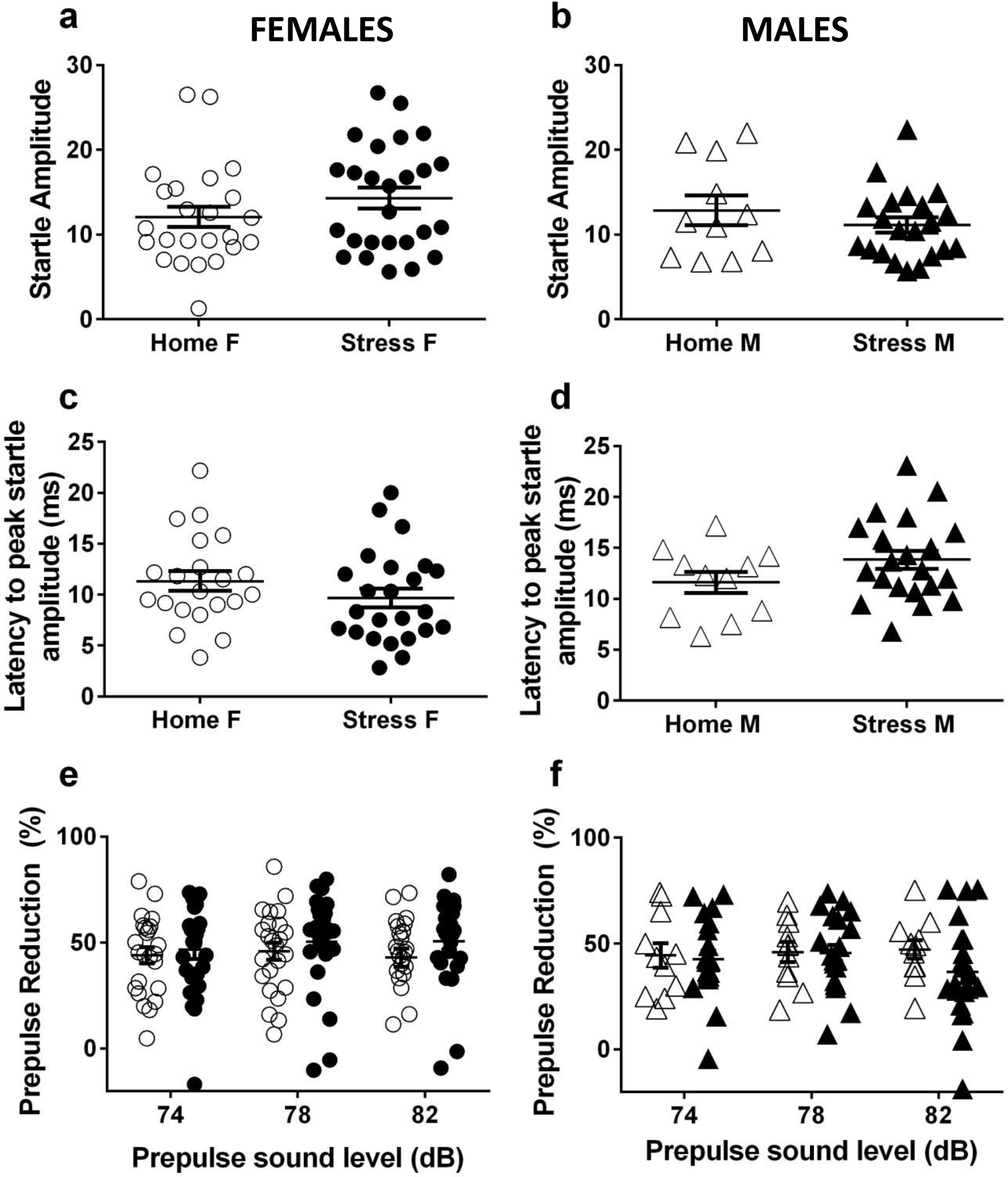
Chronic stress did not affect Acoustic Startle Response or Prepulse Inhibition in female or male mice, 2 months after stress. Both female and male mice showed no difference in startle amplitude **(a,b)**, latency to startle peak amplitude **(c,d)** or prepulse inhibition of the startle response **(e,f)** between chronically stressed and homecage control mice. (Home F: n=24; Stress F: n=26; Home M: n=11; Stress M: n=21). Error bars represent SEM.

**Table. S1.** Fear Conditioning: Significant difference in freezing were observed only in chronically stressed male mice during the Cue test. Stressed mice were compared with same-sex homecage control groups.

|  | Females |  |  |  |  | Males |  |  |  |  |
| --- | --- | --- | --- | --- | --- | --- | --- | --- | --- | --- |
| Test phase | Stress N | HC N | T stat | DF | P value | Stress N | HC N | T stat | DF | P value |
| Conditioning | 26 | 24 | 0.24 | 48 | 0.80 | 12 | 11 | 0.88 | 21 | 0.38 |
| Extinction | 18 | 21 | 0.57 | 37 | 0.56 | 12 | 11 | 0.25 | 21 | 0.80 |
| Cue Test | 26 | 24 | 0.53 | 48 | 0.59 | 12 | 11 | 2.09 | 21 | 0.04 |
| Context Test | 26 | 24 | 0.32 | 48 | 0.74 | 12 | 11 | 0.69 | 21 | 0.49 |

